# Assessment of autism zebrafish mutant models using a high-throughput larval phenotyping platform

**DOI:** 10.1101/2020.07.23.217273

**Authors:** Alexandra Colón-Rodríguez, José M. Uribe-Salazar, KaeChandra B. Weyenberg, Aditya Sriram, Alejandra Quezada, Gulhan Kaya, Emily Jao, Brittany Radke, Pamela J. Lein, Megan Y. Dennis

**Author notes:** Corresponding author: Megan Y. Dennis, Ph.D., University of California, Davis, School of Medicine, One Shields Avenue, Genome Center, 4303 GBSF, Davis, CA 95616, United States.

## Abstract

In recent years zebrafish have become commonly used as a model for studying human traits and disorders. Their small size, high fecundity, and rapid development allow for more high-throughput experiments compared to other vertebrate models. Given that zebrafish share >70% gene homologs with humans and their genomes can be readily edited using highly efficient CRISPR methods, we are now able to rapidly generate mutations impacting practically any gene of interest. Unfortunately, our ability to phenotype mutant larvae has not kept pace. To address this challenge, we have developed a protocol that obtains multiple phenotypic measurements from individual zebrafish larvae in an automated and parallel fashion, including morphological features (i.e., body length, eye area, and head size) and movement/behavior. By assaying wild-type zebrafish in a variety of conditions, we determined optimal parameters that avoid significant developmental defects or physical damage; these include morphological imaging of larvae at two time points (3 days post fertilization (dpf) and 5 dpf) coupled with motion tracking of behavior at 5 dpf. As a proof-of-principle, we tested our approach on two novel CRISPR-generated mutant zebrafish lines carrying predicted null-alleles of *syngap1b* and *slc7a5*, orthologs to two human genes implicated in autism-spectrum disorder, intellectual disability, and epilepsy. Using our optimized high-throughput phenotyping protocol, we recapitulated previously published results from mouse and zebrafish models of these candidate genes. In summary, we describe a rapid parallel pipeline to characterize morphological and behavioral features of individual larvae in a robust and consistent fashion, thereby improving our ability to better identify genes important in human traits and disorders.

**AUTHOR SUMMARY:** Zebrafish (*Danio rerio*) are a well-established model organism for the study of neurodevelopmental disorders. Due to their small size, fast reproduction, and genetic homology with humans, zebrafish have been widely used for characterizing and screening candidate genes for many disorders, including autism-spectrum disorder, intellectual disability, and epilepsy. Although several studies have described the use of high-throughput morphological and behavioral assays, few combine multiple assays in a single zebrafish larva. Here, we optimized a platform to characterize morphometric features at two developmental time points in addition to behavioral traits of zebrafish larvae. We then used this approach to characterize two autism candidate genes (*SYNGAP1* and *SLC7A5*) in two CRISPR-generated zebrafish null mutant models we developed in house. These data recapitulate previously published results related to enhanced seizure activity, while identifying additional defects not previously reported. We propose that our phenotyping platform represents a feasible method for maximizing the use of single zebrafish larvae in the characterization of additional mutants relevant to neurodevelopmental disorders.

## INTRODUCTION

Zebrafish (*Danio rerio*) are freshwater teleost fish widely used to study genes important in human diseases and traits, including neurological disorders such as autism-spectrum disorder (ASD; see review from [1]), intellectual disability (ID), and epilepsy (see review from [2,3]). Zebrafish represent a robust model for disease studies with significant genetic homology (~70-80% of orthologous genes) with humans [4,5]. Furthermore, their small size, robust reproduction, and transparent appearance during embryonic development [6] facilitate more rapid and higher-throughput experiments compared to other commonly used model organisms such as rodents. The highly efficient nature of CRISPR editing applied to zebrafish [7–9] has led to an exponential increase in the numbers of genes that can be functionally tested via direct perturbation in embryos followed by phenotypic scoring. Studies can be performed in G0 mosaic fish due to the efficient bi-allelic disruption and phenotype presentation [7,10,11]. Due to the high level of mosaicism, stable lines are readily generated allowing direct genotype-to-phenotype comparisons [12]. These approaches have successfully been applied to multiple genes in parallel in order to study their impact on neurodevelopmental features in zebrafish [11,13]. The most recent example utilized CRISPR to characterize 132 human schizophrenia gene orthologs in stable mutant zebrafish lines followed by assessment of brain morphometry, neuronal activity, and behavior, demonstrating the potential for large-scale studies of gene functions in morphology and behavior [13]. Although significant, this landmark study required considerable resources to house, cross, and phenotype individual stable mutant lines on such a large scale.

With the ease of CRISPR gene editing, our main bottleneck in analyses is quick, quantitative, and high-throughput phenotyping approaches. Currently there are few automated imaging systems for zebrafish larvae. The Vertebrate Automated Screen Technology BioImager platform (VAST BioImager™) uses a capillary-based flow system to load and image 2–7 day post fertilization (dpf) zebrafish larvae coupled with 360° rotation [14,15]. Characterization of the VAST system has demonstrated that larvae are not affected physically or developmentally by passage through the system tubing during imaging [16]. Furthermore, a number of systems have utilized successful motion monitoring of behavior of up to 96 larvae in parallel housed in plate wells, including DanioVision video tracking coupled with EthoVision software [17]. Although VAST and DanioVision provide consistent results [15], previous literature has not evaluated the combined use of VAST and a behavioral-tracking system to phenotype zebrafish mutants.

Here, we developed a high-throughput phenotyping protocol to maximize the use of multiple quantitative measurements of the same zebrafish at two developmental time points using automated imaging (VAST) and behavioral assays (DanioVision) to test for gross morphological defects and seizure susceptibility, respectively. To optimize our approach, we performed baseline experiments on a well characterized wild-type (WT) zebrafish line, NHGRI-1 [18], and subsequently tested our strategy using a novel zebrafish null model of ASD-candidate gene *SYNGAP1B*. Morpholino knockdown of the zebrafish ortholog *syngap1b* was previously shown to lead to sporadic seizures, reduced brain ventricle size, and microcephaly [19]. Mutations in orthologs of *syngap1b* in humans and rodents lead to ASD morphological phenotypes, including seizures and microcephaly [19,20]. We also tested our system using a novel zebrafish null mutant model of *SLC7A5*, a gene implicated in ASD, microcephaly, and epilepsy in humans with neurodevelopmental phenotypes also observed in mice [21]. Overall, we found that our combined phenotyping platform was capable of detecting enhanced drug-induced seizure behavior in both mutant models in addition to developmental abnormalities. Coupled with CRISPR gene editing, higher-throughput screening approaches will be instrumental in scaling up functional assessment of genes important in neurodevelopment using zebrafish.

## RESULTS

### High-throughput phenotyping of WT zebrafish larvae

#### Morphometric phenotypes

To optimize anatomical quantifications of WT (NHGRI-1) zebrafish larvae at 3 dpf (n=100) and 5 dpf (n=78) using the VAST imaging platform, we performed manual measurements of eye area (EA), body length (L), as well the width of the head at two sites previously shown to correlate with the telencephalon (Tel) and optic tectum (OT) [22] (Figure 1A). Previous work has demonstrated that larval development is not affected by passage through the VAST system tubing [16]. Using the same images, we also extracted measurements using automated analysis software FishInspector [15] for EA, L, and head width (defined as the widest part of the fish; see Materials and Methods), as well as two additional features, pericardium area, and yolk area (Figure S1A). Examining the normal distribution of data for a subset of features (EA and L), we flagged a proportion of larvae as outliers (3 dpf: 10%, 5 dpf: 7.7%; Figure S1B). By visual inspection of all images, we found that less than half of outliers represented technical errors in imaging using the VAST (e.g., full length of fish not included in the image; 3 dpf: 3%, 5 dpf: 4%) while the remaining outliers represented FishInspector software errors in automatically defining morphological features. Taking a closer look at additional measurements, including yolk and pericardium areas, as well as head width, we found that >10% of values also fell outside of the confidence interval based on a normal distribution, with the majority due to inaccuracies in FishInspector feature tracing. As such, we manually corrected FishInspector traces for EA, L, and head width, but chose not to move forward with yolk and pericardium area measurements due to difficulties in manual manipulations of these features in the software. Using this curated dataset, we showed significant correlations between our original manual measurements of EA (3 dpf: r_(94)_=0.69, *p*=1.1×10^-14^; 5 dpf: r_(72)_=0.45, *p*=5.7×10^-5^) and L (3 dpf: r_(95)_=0.69, *p* =3.3×10^-15^; 5 dpf: r_(72)_=0.96, *p* < 2.2×10^-16^) with our corrected automated measurements (Figure 1B). Additionally, we found that our manual measures of Tel correlated better to automated measures of head width at 5 dpf (3 dpf: r_(95)_= 0.13, *p*=0.196; 5 dpf: r_(71)_= 0.30, *p*=0.0087), while OT correlated with head width at both ages (3 dpf: r_(95)_=0.46, *p*=1.6×10^-6^; 5 dpf: r_(72)_= 0.35, *p*=0.0024). Based on these results, we proceeded with quantifying VAST images via FishInspector automated measurements for EA and L and manual measurements of Tel and OT as a proxy for brain size (Figure 1C).

**Figure 1.**
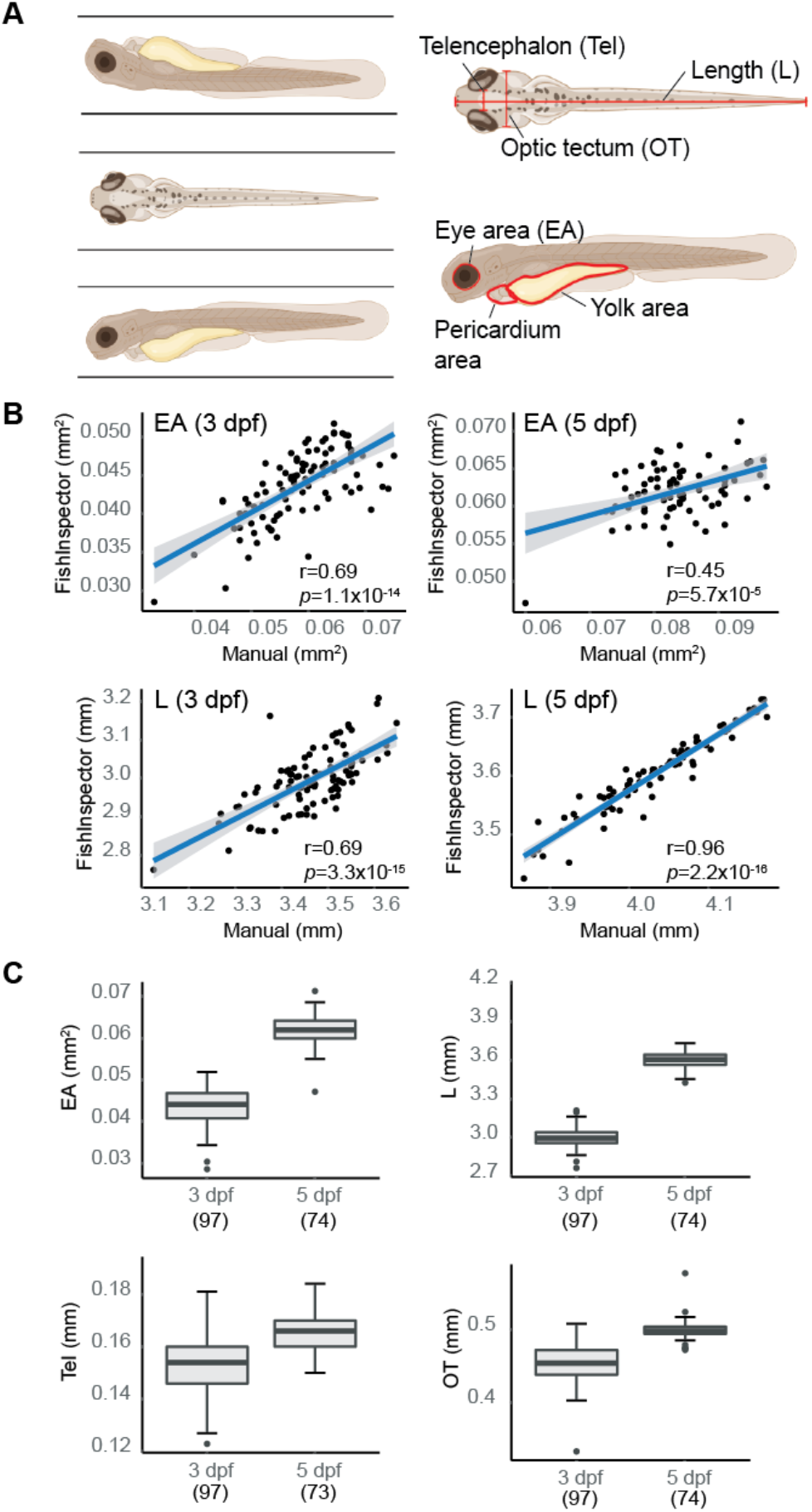
Morphometric-phenotype assessment of larvae at 3 dpf and 5 dpf. **(A)** Larval fish were imaged in three orientations (dorsal, right lateral, and left lateral) using the VAST system. Morphometric features were obtained manually using Fiji (L, EA, Tel, and OT) and automatically using FishInspector (L, EA, head width, pericardium area, and yolk area). **(B)** Manual and automated measurements were compared with L and EA (averaged between both eyes) showing high correlations at 3 dpf (n=97) and 5 dpf (n=73) (Pearson correlation). Blue line represents the fit of a linear model with 95% confidence intervals. **(C)** Boxplots pictured represent distribution of measurements for larval morphometric features obtained via automated (L and EA) and manual (Tel and OT) methods. Boxplots include the median value (dark line) and the whiskers represent a 1.5 interquartile range.

#### Behavioral phenotypes

A number of studies have used motion tracking to characterize locomotor behavior and seizure susceptibility in order to assay neurological phenotypes in zebrafish larvae [19,23,24]. Previous work using the chemical convulsant pentylenetetrazol (PTZ) has identified a concentration dependent difference in distance, velocity, and swim activity [25,26]. To verify our ability to recapitulate these results, we subjected larvae (5 dpf) to varying concentrations of two GABAA receptor antagonists, PTZ (2 mM, 10 mM, 15 mM; n=12), a selective antagonist, as well as bicuculline (BCC; 0 μM, 5 μM, 10 μM; n=10), a non-selective antagonist. We then tracked their movement over one hour (h) with a brief flashing of lights after 45 minutes (min) using the DanioVision motion-tracking system (Figure 2). We quantified the average distance of larval movement per min over the entire 1 h and over the last 15 min after being subjected to flashing light, respectively. For maximum concentrations of BCC (10 μM) and PTZ (15 mM), we observed reduced movement overall versus lower concentrations, suggesting larvae experienced what is known as stage III clonal seizures, consistent with has been previously shown with 15 mM PTZ [25]. Overall, we found the strongest effect on distance moved (mean distance=166.88 mm) with the lowest variance (SD=27.17) using 10 mM PTZ, a concentration with little morbidity, in concordance with previous studies [23,25].

**Figure 2.**
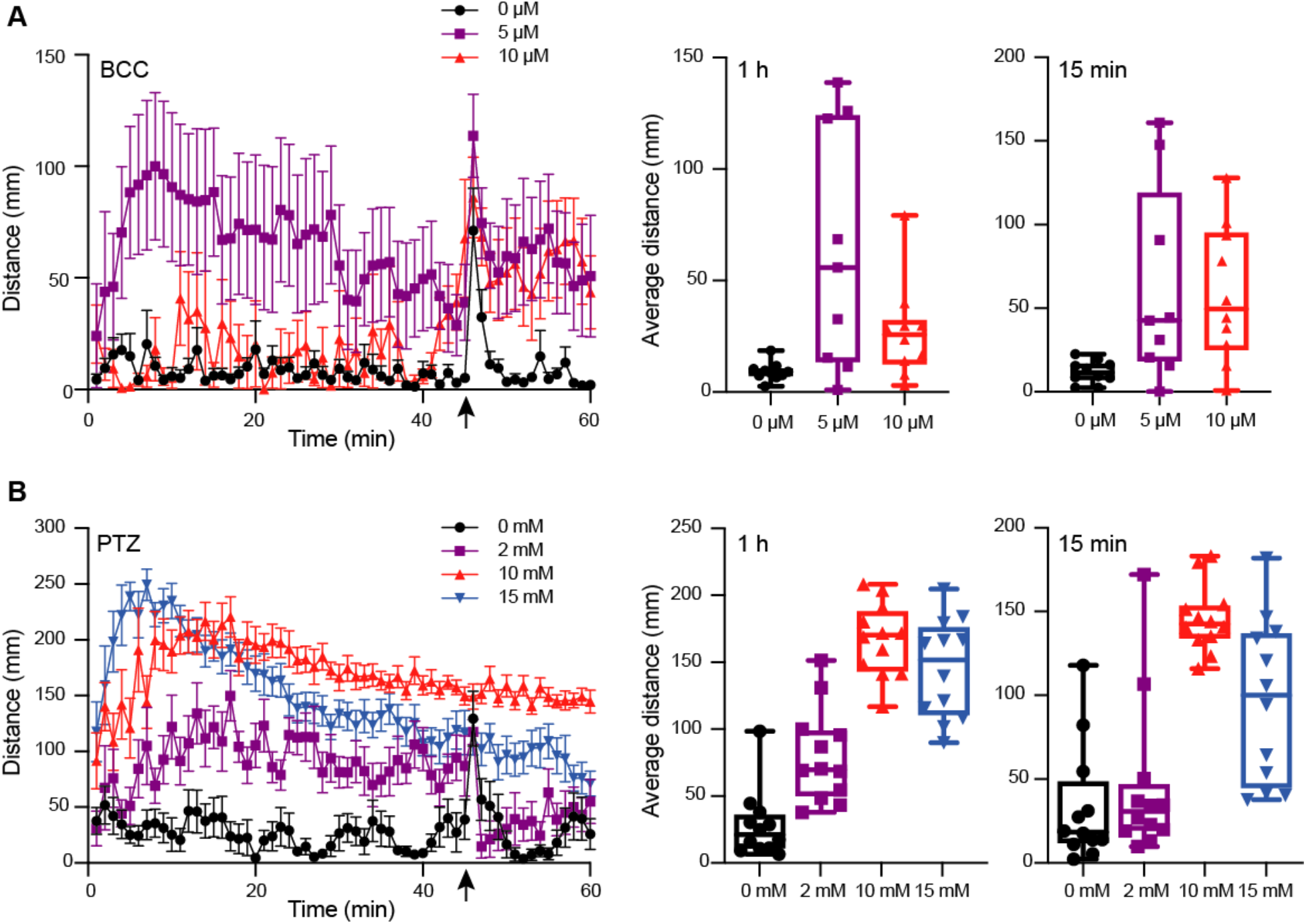
Behavioral phenotype assessment of larvae at 5 dpf using motion tracking. Larvae treated with varied concentrations of GABAA antagonizing drugs (**A)** BCC (0 to 10 μM; n=10) or (**B**) PTZ (0 to 15 mM; n=12), respectively, were tracked using DanioVision and total distance moved per minute measured for one hour. With 15 min remaining (indicated by an arrow), larvae were subjected to flashing lights for one min. The average movement per fish across 60 min and in the last 15 min following flashing lights are indicated as boxplots for each drug concentration. Left plots include the mean distance per minute and with error bars representing standard deviation values.

### Optimization of a combined morphological and behavioral phenotyping platform

Although previous studies have shown the utility of VAST and DanioVision platforms to independently characterize phenotypes in a high-throughput fashion, we sought to identify optimal parameters to utilize these approaches in combination at multiple developmental time points in individual zebrafish larvae while also minimizing impacts on traits. WT NHGRI-1 embryos (n= 992) collected across seven experiments were placed in 96-well plates and subjected to varied measurements/treatments, including VAST imaging at 3 dpf, behavioral tracking at 5 dpf using the DanioVision with (10 mM) or without (0 mM) PTZ, and VAST imaging at 5 dpf (Figures 3A and S2, Table S1). Using our combined dataset, we found that larvae subjected to VAST at 3 dpf exhibited smaller measurements at 5 dpf for EA (15.2% decrease, *p*=4.85×10^-4^), Tel (13.6% decrease, *p*=1.36×10^-2^), and OT (4.5% decrease, *p*=7.77×10^-4^), in addition to a notable decrease in the average movement observed during the behavioral assays (0 mM PTZ; average movement 1 h: 47.7% decrease, *p*=0.046, average movement last 15 min: 60.9% decrease, *p* = 0.017) (Table 1, Figures 2A and S3). Treating larvae with PTZ during our behavioral screen also had an effect on morphometric measurements at 5 dpf, with larvae exhibiting a reduction of 4.3% in EA (*p*=8.77×10^-3^), 3.3% in L (*p*=3.89×10^-9^), 10.2% in Tel (*p*=2.28×10^-3^), and 3.1% in OT (*p*=5.53×10^-4^). Additionally, we observed an increase in mortality (~4%) for PTZ-treated versus untreated larvae. As such, we moved forward with a final phenotyping platform that included VAST imaging at 3 dpf followed by motion tracking at 5 dpf with half of the larvae treated with (10 mM) or without (0 mM) PTZ. To minimize PTZ effects on morphometric measurements and mortality, we performed final VAST imaging at 5 dpf only in larvae not treated with the drug (Figure 3B). We determined that ~50 fish are needed per genotype in order to detect a >4% change between groups for morphometric features at 5 dpf at 80% power using our ‘optimized’ assay, though for certain traits (OT and Tel) far fewer fish are required (Table S2, Figure S4).

**Figure 3.**
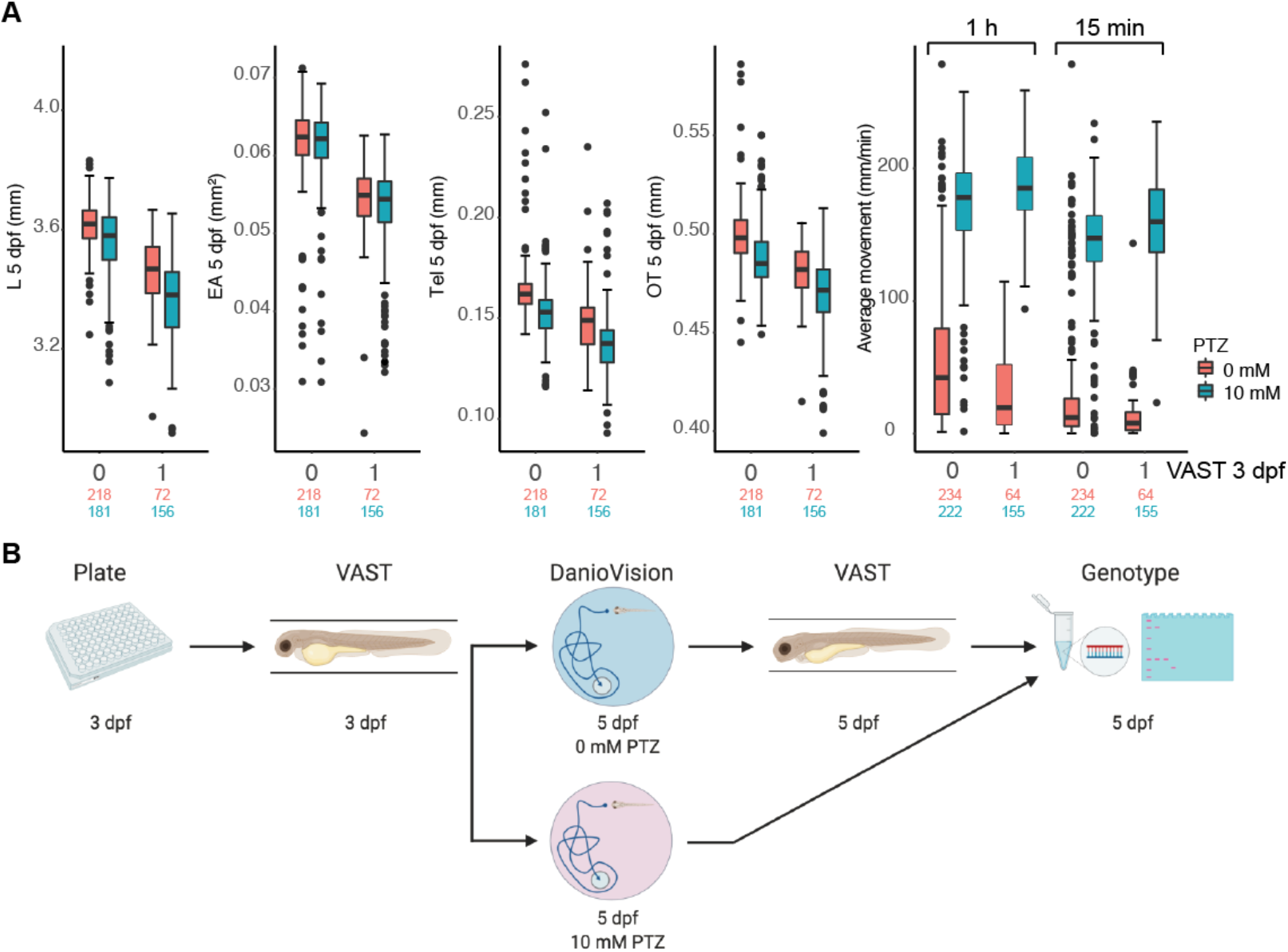
High-throughput zebrafish larvae phenotyping platform. **(A)** Impact of morphometric (L, EA, Tel, OT) and behavioral (average movement for the entire one h and the final 15 min following flashing lights) measurements for 5 dpf larvae for fish not subjected (0) and subjected (1) to VAST measurements at 3 dpf are shown as boxplots. Behavior was also plotted at varied PTZ concentrations (red=0 mM and blue=10 mM). Total numbers of measured larvae are indicated in parentheses. Boxplots for morphological and behavioral measurements include the median value (dark line) and the whiskers represent a 1.5 interquartile range. **(B)** The final combinatorial phenotyping platform is pictured, which combines two morphometric measurements (3 and 5 dpf) and behavioral tracking (5 dpf) with 0 mM and 10 mM PTZ, followed by genotyping of larvae.

**Table 1.** Summary of effect size and significance for each treatment per measurement.

| Morphometric measurement<br>(5 dpf) | Complete model* |  |  | VAST at 3 dpf |  | Behavior |  | PTZ |  |
| --- | --- | --- | --- | --- | --- | --- | --- | --- | --- |
|  | F (df) | p-value | R<sup>2</sup> | Beta | p-value | Beta | p-value | Beta | p-value |
| Average EA<br>(mm<sup>2</sup>) | 43.98<br>(9, 617) | p < 2.2x10<sup>-16</sup> | 0.382 | -<br>0.00553 | 4.85x10<sup>-4</sup> | 0.0006 | 0.477 | -0.0020 | 8.77 x 10<sup>-3</sup> |
|  |  |  |  | Yes= 228, No= 399 |  | Yes= 566, No= 61 |  | 10mM= 337, 0mM= 290 |  |
| L<br>(mm) | 69.98<br>(9, 617) | p < 2.2x10<sup>-16</sup> | 0.498 | -<br>0.06075 | 0.0525 | 0.0316 | 0.068 | -0.0923 | 3.89x10<sup>-9</sup> |
|  |  |  |  | Yes= 228, No= 399 |  | Yes= 566, No= 61 |  | 10mM= 337, 0mM= 290 |  |
| Tel<br>(mm) | 35.71<br>(9, 617) | p < 2.2 x10<sup>-16</sup> | 0.333 | -<br>0.01152 | 0.0136 | -<br>0.0053<br>86 | 0.037 | -0.0080 | 5.53x10<sup>-4</sup> |
|  |  |  |  | Yes= 228, No= 399 |  | Yes= 566, No= 61 |  | 10mM= 337, 0mM= 290 |  |
| OT<br>(mm) | 39.84<br>(9, 615) | p < 2.2 x10<sup>-16</sup> | 0.359 | -<br>0.01583 | 7.77x10<sup>-4</sup> | -<br>0.0004 | 0.866 | -0.0071 | 2.28x10<sup>-3</sup> |
|  |  |  |  | Yes= 226, No= 399 |  | Yes= 564, No= 61 |  | 10mM= 335, 0mM= 290 |  |
| 1 h average distance (mm) | 186.5<br>(8, 666) | p < 2.2 x10<sup>-16</sup> | 0.688 | 9.916 | 0.046 | - | - | 119.6 | < 2x10<sup>-16</sup> |
|  |  |  |  | Yes= 219, No= 456 |  | - |  | 10mM= 377, 0mM= 298 |  |
| 15 min average distance (mm) | 184.3<br>(8, 666) | p < 2.2 x10<sup>-16</sup> | 0.685 | 11.712 | 0.017 | - | - | 122.7 | < 2x10<sup>-16</sup> |
|  |  |  |  | Yes= 219, No= 456 |  | - |  | 10 mM= 377, 0 mM= 298 |  |

\* The 'complete model' also included 'Experiment' as a variable to reduce inter-batch effects (see Methods).

### Proof of concept using a stable zebrafish mutant model of *SYNGAP1*

Though the platform itself significantly impacts morphometric and behavioral features in WT larvae, we hypothesized that effects would be equal across all fish, thus making it possible to parse genotype effects on development and behavior in mutant larvae. To test this, we used CRISPR to generate a null mutant zebrafish model of human *SYNGAP1*, a gene encoding neuronal Ras and Rap GTPase-activating proteins. Heterozygous mutations of *SYNGAP1*, important for synaptogenesis and regulation of excitatory synapses [27,28], are associated with neurodevelopmental disorders, including epilepsy, ASD, and ID [29]. A previous study using morpholinos targeting mRNAs encoding the two zebrafish orthologs of the gene (Figure S5A) *syngap1a* and *syngap1b* showed malformed mid- and hindbrain (48 hours post fertilization (hpf)), developmental delay (48 hpf), and spontaneous seizures, correlated to an observed decrease in GABA neurons in the mid- and hindbrain, specific to the *syngap1b* morphant larvae at 3 dpf [19]. Although functional domains from *syngap1a* and *syngap1b* are conserved in zebrafish, the characterization of *syngap1a* was not performed in the aforementioned study due to low penetrance of the morpholino and toxicity of using higher doses [19]. The longitudinal expression profile of *syngap1b* is relatively low until 30 hpf based on previously-published RNA-seq data [30] (Figure S5B), suggesting that morpholinos, which are best suited for knocking down early developmental genes, may not represent an optimal approach to characterize *syngap1b*. As such, we generated a stable mutant line *syngap1b^tup1^* carrying a 23-bp frame-shifting deletion predicted to produce a transcript encoding a 213-amino acid protein (84.12% shorter than the full-length protein) (Figure 4A). To ensure any phenotypes produced in *syngap1b^tup1^* were not a product of off-target mutations, we also assayed the top 13 genetic loci predicted by CIRCLE-seq [31] to be edited and detected no indels (Table S3, Figure S5C), including in the *syngap1a* paralog.

**Figure 4.**
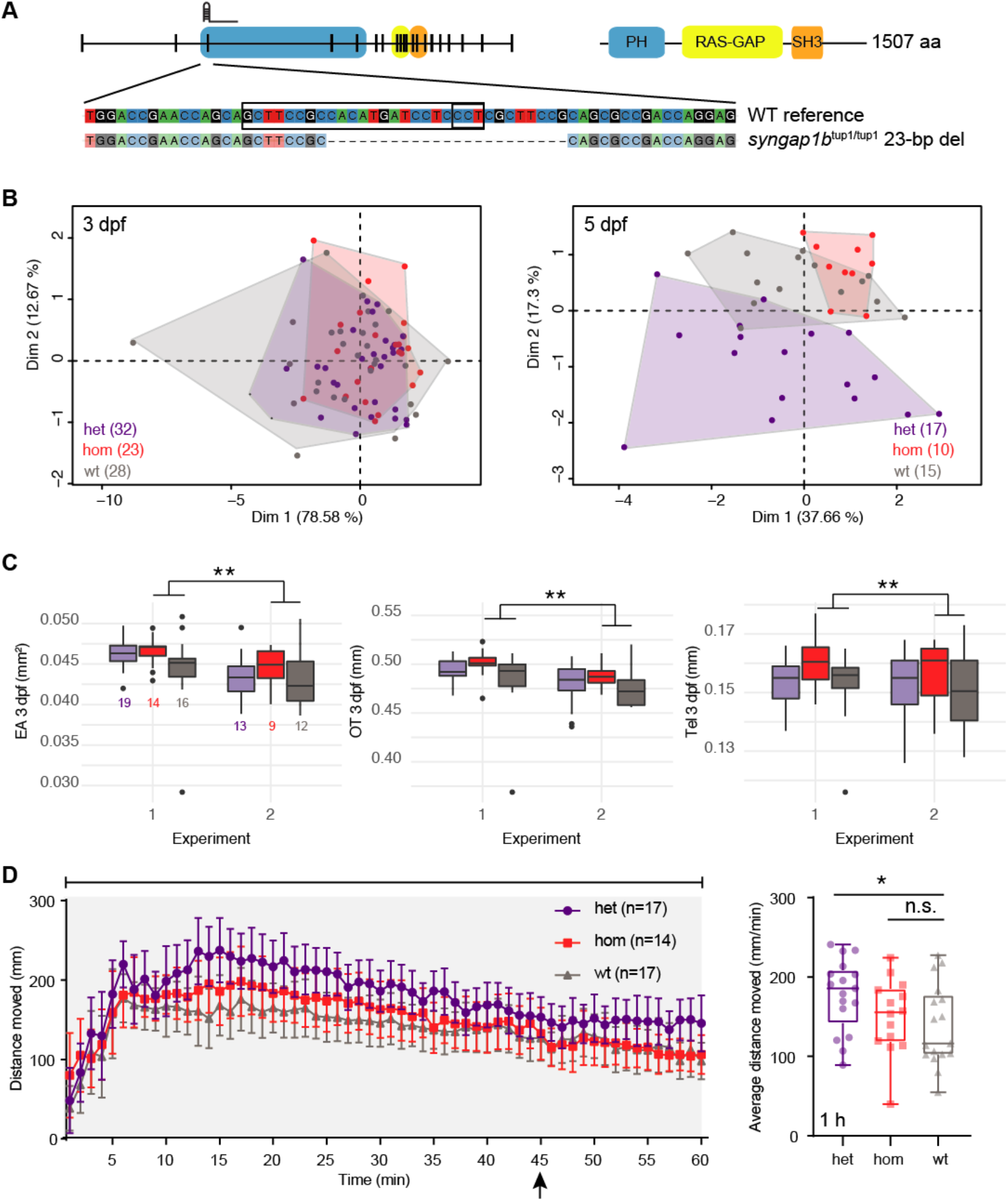
*syngap1b* CRISPR mutants exhibit developmental defects and enhanced seizures. **(A)** The zebrafish mutant line *syngap1b*^tup1/tup1^ was generated using a single gRNA targeting the third exon of the gene resulting in a 23-bp deletion leading to a frameshift. **(B)** PCA of combined morphometric traits at 3 and 5 dpf colored by genotype. **(C)** The three traits found to be significantly increased in het (purple) and hom (red) mutant versus WT (gray) siblings included EA, OT, and Tel at 3 dpf, displayed as box plots with total numbers of larvae measured indicated in the EA plot (**, *p* < 0.01 using a Tukey post-hoc test). Boxplots include the median value (dark line) and the whiskers represent a 1.5 interquartile range. **(D)** Behavioral data from DanioVision motion tracking shows significant increased distance moved over the entire 1 h for het versus WT siblings (*, *p* < 0.05 using a Dunn’s multiple comparison post-hoc test). The left plot indicates the mean distance per minute with error bars representing standard deviation. The box plot shows the mean for the 1 hr behavioral assay, the median value (dark line) and the whiskers represent a 1.5 interquartile range.

We collected data on siblings (n=124) produced by crossing *syngap1b^tup1+^* lines, creating a mix of WT, heterozygous (het), and homozygous (hom) larvae, using our multi-phenotyping platform (Table S4). We observed a skew in genotype frequency (n=48 het, n=33 hom, and n=43 WT; *p*=0.02 *χ*^2^ test) from the expected Mendelian distribution suggesting that *syngap1b^tup1^* mutants may be less viable compared to their WT siblings. We did not observe an enrichment of larvae carrying mutant alleles falling outside of a normal distribution of morphometric measurements, ruling out highly-penetrant developmental defects associated with null mutations of *syngap1b* (Figure S5D). Considering all morphometric data in combination, we performed a principal-component analysis (PCA) and identified that L, EA, OT, and Tel all contributed significantly to the dispersion of our data (all squared loadings > 0.60 for first dimension) (Table S5, Figure 4B). Therefore, we analyzed our morphometric data using models that included each measurement as a covariate to identify individual effects of *syngap1b* genotype. This combined approach revealed larger EA (4.6% increase, *p*=4.16×10^-5^ Tukey post-hoc), OT (3.1% increase, *p*=2.7×10^-6^ Tukey post-hoc), and Tel (4.1% increase, Tukey post-hoc: *p*=0.016) in hom *syngap1b^tup1/tup1^* larvae at 3 dpf relative to their WT siblings (Table S5, Figure 4C). EA was also suggestive to be larger on hom *syngap1b^tup1/tup1^* at 5 dpf (*p* = 0.053 Tukey post-hoc). Notably, hom *syngap1b^tup1/tup1^* larvae did not exhibit increased L (3 dpf: *p*=0.267; 5 dpf: *p*=0.348 Tukey post-hoc) suggesting that mutant larvae were not generally larger overall. Next we assessed seizure susceptibility of larvae when treated with PTZ and found a significant increase in movement for het *syngap1b^tup1/+^* mutants (mean distance=177.38 mm, *p*=0.047 Dunn’s multiple comparison post-hoc) compared with their WT siblings (mean distance=138.50 mm) (Table S6, Figure 4D). Notably, no significant difference was observed for hom *syngap1b^tup1/tup1^* mutants (mean distance=150.56 mm), nor in larvae not treated with PTZ (Figure S5E).

### A novel zebrafish model for ASD-candidate gene *SLC7A5*

Recently, recessive mutations of *SLC7A5* were reported in children with ASD, microcephaly, and seizures [21]. This gene encodes large neutral-amino-acid transporter 1 (LAT1), an essential channel present at the blood brain barrier in epithelial cells, that facilitates the movement of branched-chain amino acids and other large neutral amino acids from the periphery into the brain [32]. Knockout mouse models recapitulated several phenotypic features observed in patients, including ASD-related behaviors, motor coordination abnormalities, and alterations in inhibitory and excitatory neuronal communication. We tested if ‘knockout’ of the single zebrafish ortholog *slc7a5* resulted in similar neurodevelopmental defects (Figure S6A). Using published RNA-seq whole-embryo data [30], we found that *slc7a5* is expressed starting between 6 and 24 hpf followed by a steady increase throughout larval development (Figure S6B). To model a complete homozygous recessive knockout, we generated the stable mutant line *slc7a5^tup1^* carrying an 18-bp deletion predicted to encode a truncated protein of 17 amino acids (96.75% shorter than the full-length protein) (Figure 5A). We verified that no indels existed at the top six off-target sites impacting genes as predicted by CIRCLE-seq that might contribute to phenotypes (Table S3, Figure S6C).

**Figure 5.**
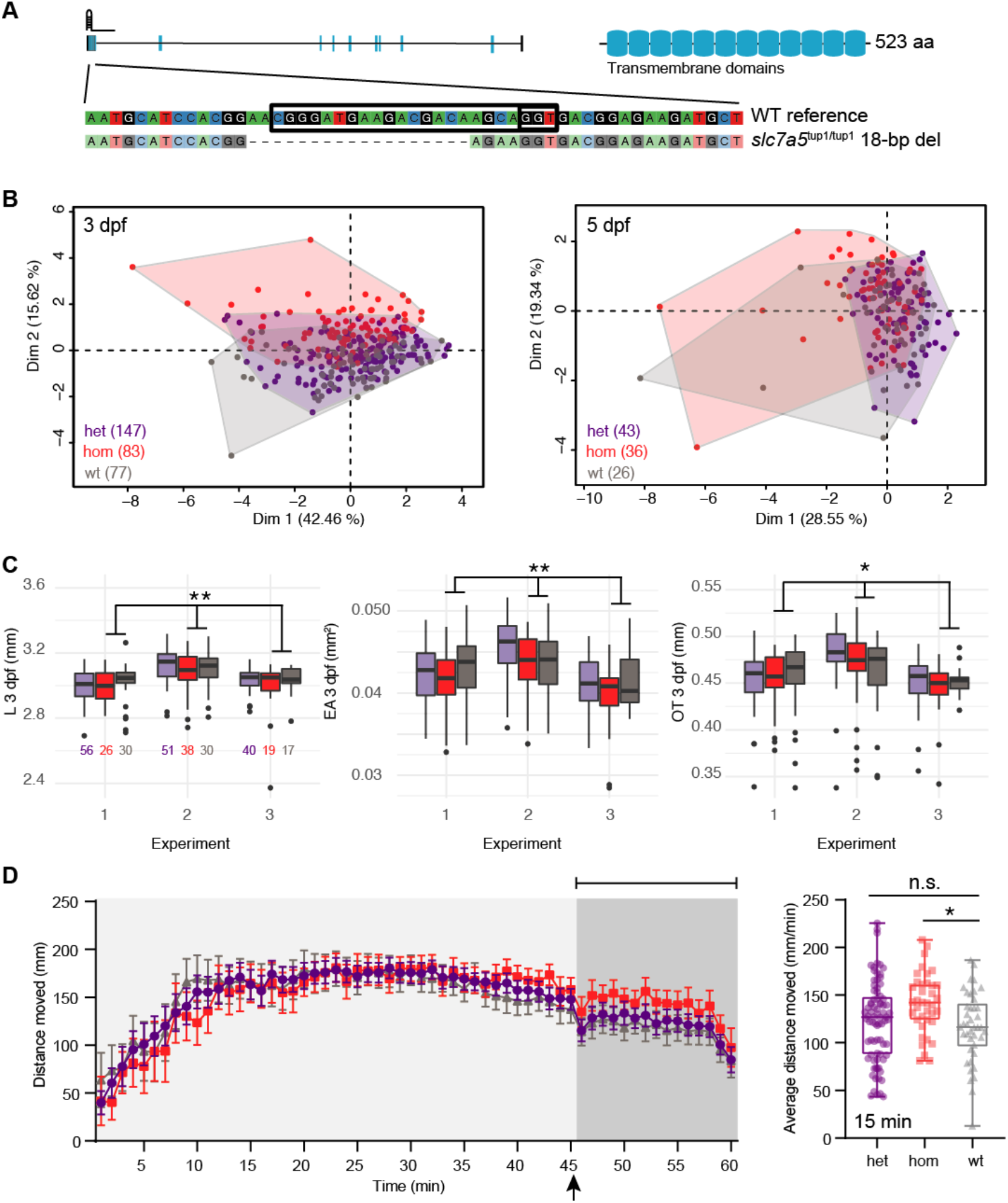
*slc7a5* CRISPR homozygous mutants exhibit moderate developmental delay and enhanced seizures with flashing lights. **(A)** The zebrafish mutant line *slc7a5*^tup1/tup1^ was generated using a single gRNA targeting the first exon of the gene resulting in an 18-bp deletion leading to a frameshift. **(B)** PCA of combined morphometric traits at 3 and 5 dpf colored by genotype. **(C)** The three traits found to be significantly increased in het (purple) and hom (red) mutant versus WT (gray) siblings included L, EA, and OT at 3 dpf, displayed as box plots with total numbers of larvae measured indicated in the EA plot (*, *p* < 0.05; **, *p* < 0.01 using a Tukey post-hoc test). Boxplots include the median value (dark line) and the whiskers represent a 1.5 interquartile range. **(D)** Behavioral data from DanioVision motion tracking shows significant increased distance moved over the final 15 min following flashing lights for hom versus WT siblings (*, *p* < 0.05 using a Dunn’s multiple comparison post-hoc test). The left plot indicates the mean distance per minute with error bars represented as standard deviation. The box plot shows the mean for the final 15 min of the behavioral assay, the median value (dark line) and the whiskers represent a 1.5 interquartile range.

Using data collected from siblings (n=337) produced from heterozygous G_1_ crosses, we did not observe any significant mortality nor unexpected skew in genotypes (n=161 het, n=90 hom, and n=86 WT; *p*=0.81 *χ*^2^ test) showing that *slc7a5^tup1^* mutants are viable compared to their WT siblings (Table S7). Examining the distribution of morphometric data, we also did not note any enrichment of mutants in fish exhibiting outlier traits (Figure S6D). When considering all features in combination, we noted that hom *slc7a5^tup1/tup1^* mutants clustered separately from the other two genotypes at 3 dpf (Figure 5B). The primary traits driving this deviation were EA, OT, and L (all squared loadings > 0.80 for first dimension). In models that accounted for these multiple variables simultaneously, we observed that hom *slc7a5^tup1/tup1^* exhibited significant, albeit modest, differences in EA (0.9% decrease, *p*=7.5×10^-3^ Tukey post-hoc), OT (0.1% increase, *p*=0.023 Tukey post-hoc), and L (0.6% decrease, *p*=9.9×10^-3^ Tukey post-hoc) at 3 dpf (Table S5, Figure 5C). The overall reduction of eye area, head size, and body length suggests larvae may experience minor developmental delay. At 5 dpf, no measurement showed differences between *slc7a5* genotypes. Furthermore, we quantified significantly greater movement in PTZ-treated larvae following light flashes in the final 15 min, suggesting enhanced seizures in hom *slc7a5^tup1/1up1^* mutants (mean distance=141.3 mm, *p*=0.013 Dunn’s multiple comparison post-hoc) versus WT (mean distance =116.7 mm) that was not detected for het siblings (Table S6, Figure 5D). Both of these observations are in line with the recessive nature of *SLC7A*5 mutations associated with defects identified in children with ASD. Notably, we did not identify any significant differences across genotypes when considering distance moved over the entire 1 h nor for larvae not subjected to PTZ (Figure S6D).

## DISCUSSION

Several studies have proposed morphological and behavioral phenotyping platforms for the characterization of zebrafish larvae [14,15,33,34]. However, previous literature has not assessed the impact of combining automated VAST morphometric and DanioVision behavioral assays in the same larvae, and particularly for the purpose of testing CRISPR mutants. Here, our goal was to determine if imaging with VAST or motion tracking would alter zebrafish behavior or developmental features, in addition to optimizing parameters to maximize the number of measurements obtained for each larva. Contrary to a previous study [14], we found that imaging 3 dpf larvae using VAST significantly impacted downstream behavior and development measured at 5 dpf. Despite the impacts on downstream developmental features, the benefits of using the VAST platform are significant, as it facilitates automated measurements in larvae while still alive, thus preventing manual errors, improving consistency, and increasing speed of measurements. One significant downside is that larvae must be temporarily anesthetized for imaging, though the low tricaine methanesulfonate (MS-222) concentration used in our studies (76 μM or 20 mg/L) [35] would suggest that impacts on development might be minimal. Previous studies have shown that focal exposure to varying concentrations of MS-222 (191–574 μM) at earlier developmental time points (before 26 hpf) leads to cartilage malformations [36] and upregulation of genes associated with apoptosis [37]. There is a considerable body of literature demonstrating the negative effects of anesthesia in the developing human brain [38], with most studies finding associations with neuronal apoptosis specifically when using anesthetics. More relevant to our study is the cell death by necrosis and apoptosis observed when using local anesthetics, which act in voltage gated sodium channels, as is the case with MS-222 [39]. However, the effects of MS-222, an anesthetic used in fish not humans, have not been well studied in the developing nervous system of zebrafish [40]; thus, future studies are needed.

To assess whether our platform was capable of identifying defects in spite of impacts the assays had on measured features, we proceeded with characterizing two novel CRISPR-generated ASD mutant zebrafish lines (*syngap1b* and *slc7a5*). We assayed a stable mutant line of *syngap1b* as a proof of concept, a gene that had previously been studied using morpholinos [19]. We noted a skewed distribution of genotypes (reduced hom and het alleles versus WT) after subjecting them to our combined phenotyping assay, suggesting that mutant larvae have reduced viability, in line with perinatal lethality of hom null mutations observed in *Syngap1* mouse models [29]. Our het *syngap1b^tup1/+^* mutants moved greater distances over 1 h in the presence of PTZ versus WT, suggesting they may be experiencing enhanced seizures, consistent with spontaneous seizures in *syngap1b* morphants [19]. In contrast, hom *syngap1b^tup1/tup1^* mutants did not significantly differ from WT or hom mutants in their overall movement, suggesting they perhaps exhibit intermediate seizure effects (Figure 4D). Notably, our assay does not currently score for spontaneous clonic seizures observed previously in morphants but may explain the reduction in movement of our hom versus het mutants; as such, automated methods to detect such behaviors in PTZ-treated and -untreated larvae from motion-tracking videos represents a future direction. Furthermore, our morphometric data suggests that *syngap1b^tup1^* mutants exhibit significantly increased head and eye sizes at 3 dpf, counter to the findings in zebrafish morphants [19], mutant mice [41], and patients [29] that exhibit reduced brain ventricle sizes and mild microcephaly. It is important to note that microcephaly phenotypes have low penetrance in *SYNGAP1* zebrafish morphants and are identified in 30% of human patients carrying heterozygous mutations of the gene [42]. Traits with low penetrance are difficult to detect using our higher-throughput pipeline, which compares means across groups requiring a larger number of fish to detect sporadic differences.

Finally, we generated a novel stable mutant of *slc7a5*, a gene never before assayed in zebrafish, with the goal of recapitulating microcephaly and seizure phenotypes observed in human patients and mouse models carrying homozygous recessive mutations [21]. Our PCA analysis of all morphometric features at 3 dpf showed that hom *slc7a5^tup1/tup1^* mutants clustered separately from WT and het siblings, though individual traits were not significantly different between genotypes. Since our assay quantified multiple traits from the same larvae, we could parse individual effects of genotype on each measurement by considering each measured trait as a covariate. By doing this, we detected small yet significant reductions in body length, eye area, and head size (OT) in our hom *slc7a5^tup1/tup1^* mutants at 3 dpf compared to both het and WT. Furthermore, hom larvae exhibited increased movement, which could indicate enhanced seizures, in the presence of light flashing, but not over the entire one hour when treated with PTZ. Although not reported in the few *SLC7A5* patients published to date, this suggests that patients may be susceptible to light-flashing-induced seizures, as is the case with certain types of epilepsy in humans [43]. For both *slc7a5* and *syngap1b*, we did not observe any strong mutational effects in non-PTZ treated larvae. For future studies, we will consider incorporating additional perturbations during motion tracking with known associations to enhanced behaviors and seizure phenotypes, such as heat [44,45] and startle [46].

In all, we have demonstrated that our multi-phenotyping platform can detect morphological and behavioral changes in two ASD mutant zebrafish models. This approach, coupled with CRISPR mutant screens of many genes in parallel, either in G_0_ mosaic or stable mutant lines, represents a powerful first-pass to quickly identify genes impacting development and behavior. Though we focus our study here on ASD genes, most of the traits we assessed are not unique to neurodevelopment; thus, additional studies are required to pinpoint affected brain features. Future work includes expanding our phenotypes to include more neurological traits. In addition to the aforementioned improvements to our assay to induce and detect spontaneous seizures, we aim to extract more phenotypes from our VAST images, in addition to L, EA, Tel, and OT. By incorporating transgenic reporter fish, such as those with labeled excitatory and inhibitory neurons to identify neuronal transmission imbalance or calcium reporters such as GCaMP to further characterize seizures, coupled with a higher resolution fluorescent microscope and VAST imaging, our assay could more comprehensively characterize specific brain defects in our disease mutant models. Furthermore, a current limitation to measuring additional traits is the reliability of existing automated methods to accurately and consistently extract features. Although many of our measurements were performed automatically with the FishInspector software, we manually inspected every image due to issues with feature tracing to ensure accuracy and, in some cases, deferred to manual measurements (OT and Tel) or removed features completely from our analyses (HW, Y, and P). Future goals include improvements in automated extraction of features as well as integrating data from multiple sources to pinpoint genotype effects.

## MATERIALS AND METHODS

### Zebrafish care and husbandry

All experimental procedures and animal handling were performed in accordance with the University of California Davis and AAALAC guidelines and were approved by the Institutional Animal Care and Use Committee (protocol #20690; Office of Laboratory Animal Welfare Assurance Number D16-00272 (A3433-01)). Adult zebrafish from an NHGRI-1 background [18] were housed in a modular zebrafish system (Aquaneering, San Diego, CA) and all fish were kept in a 10 h dark/14 h light cycle, and 28 ± 0.5°C filtered, UV sterilized water [6]. NHGRI-1 is an isogenic line that has been fully sequenced and was used for all of our studies. Adult zebrafish were monitored and fed two times a day: flakes (Zebrafish select diet, Aquaneering, San Diego, CA) and brine shrimp (Aquaneering, San Diego, CA). For experimentation, we collected eggs from randomly assigned NHGRI-1 pairs from at least ten breeding pairs and a total of seven experiments for WT. For *syngap1b* mutants, we collected eggs from four breeding pairs over a total of two experiments. For *slc7a5* mutants, we collected eggs from five breeding pairs over a total of three experiments. For breeding, we kept our adult female and male fish separate overnight in a one liter crossing tank (Aquaneering, item #ZHCT100, San Diego, CA) and released them, by removing the divider in the crossing tank when the lights turned on the next morning. We collected the fertilized embryos one hour after the fish were released. Eggs were then rinsed two times with embryo water (0.03% Instant Ocean salt in deionized water at 28 ± 0.5°C) to remove debris and plated at a maximum of 100 embryos per plate [47]in standard (100 x 15 mm) Petri dishes in embryo water and incubated at 28 ± 0.5°C until they were used for experiments at 3 and 5 dpf. We used a Leica dissecting microscope (Buffalo Grove, IL) to quantify, stage, monitor, and remove dead embryos daily, following previously described methods [6]. Embryos were mixed from all clutches obtained (from crossed pairs) and randomly assigned to experimental groups and or CRISPR microinjections.

### Chemicals

Tricaine Methanesulfonate (MS-222, purity ≥ 99%) (Thermo Fisher Scientific, Waltham, MA) was prepared as 2x stocks in embryo water and diluted to its final concentration (0.02 mg/ml for VAST or 0.125% for adult zebrafish genotyping) the day of experiments. All zebrafish larvae that were used for morphometry experiments were exposed to MS-222 through the VAST (Union Biometrica, Holliston, MA) platform. PTZ (purity ≥ 99%, Sigma-Aldrich) and BCC (purity ≥ 97%, Sigma-Aldrich) solutions were prepared the day of experiments in embryo water and their concentrations were selected based on previous studies [25,48] and our baseline exposure experiments.

### VAST

For morphological measurements we used the VAST instrument equipped with a large particle sampler (LP sampler^™^) on fish at 3 or 5 dpf. We collected images using the VAST built-in camera and measured EA, HW, and L from single fish that were imaged ventrally and dorsally. Our parameters for VAST imaging at 3 and 5 dpf were similar with the exception of the template images, which correspond to the correct age and body length, which was 3.8 mm for 3 dpf fish and 4 mm for 5 dpf larvae, following a previously-described protocol [16]. Images were processed with FishInspector imaging software (version 1.03 [15]) [15] to extract all established shapes for each measured larvae. After this, specific morphometric measurements were extracted using the R package Momocs [49] with the known capillary width as a scale. Some images in FishInspector required manual correction performed by using the “edit” function. EA represented the average of both eyes, extracted by tracing the outside of each eye and measuring the area of this shape. L and head width were measured using the outline of the fish in the dorsal view and extracting the distance between the farthest most left and right (L) and top and bottom (head width) points of the fish, respectively. Pericardium area was measured by tracing the pericardial sac and extracting its area. Yolk area was measured by tracing the yolk, which presents as yellow in color, and extracting its area. Manual measurements were also performed for OT (the distance directly behind the eyes) and Tel (the distance directly between the eyes) following a previously described method [22]. FIJI was used for the manual measurements using a set scale, taking the width of the capillary as a reference.

### Behavioral assay

Locomotor behavior screening of zebrafish larvae in round 96-well plates was performed in WT NHGRI-1 and stable mutant lines at 5 dpf using the DanioVision (Noldus, Leesburg, VA) instrument. Zebrafish larvae were individually placed (one fish per well) in 100 μl of embryo water. Experiments were conducted for a total of 1 h. Habituation was performed for 10 minutes by incubating the 96-well plate with larvae in the DanioVision chamber (28.5°) prior to every experiment. Then 100 μl of embryo water (0 mM PTZ) or 100 μl of 4, 20, or 30 mM PTZ (final concentration of 2, 10, or 15 mM) was added for a final volume of 200 μl per well. For experiments with BCC, addition of 100 μl of embryo water (0 μM BCC) or 100 μl of 10, or 20 μM BCC (final concentration of 5 and 10 μM) was added for a final volume of 200 μl per well. The experiment started immediately following exposure in the temperature controlled (28.5°) DanioVision chamber. After 45 min of motion tracking with light, we implemented light flashes (three flashes for 10 seconds every 10 seconds) to assay light-induced seizurogenic activity [24]. Locomotor behavioral activity, quantified as total distance moved per min over 1 h, was recorded with EthoVision 10.0 tracking software.

### Zebrafish CRISPR mutant generation

Mutant zebrafish lines (*syngap1b* and *slc7a5*) were created by microinjections of ribonucleic proteins (RNPs) comprising Cas9 enzyme coupled with single guide RNAs (gRNAs) (Integrated DNA Technologies, Coralville, IA) following the manufacturer’s protocol (Integrated DNA Technologies; see Table S3 for description of gRNAs). Microinjections were performed at the one-cell stage as previously described [7], using needles from a micropipette puller (Model P-97, Sutter Instruments, Novato, CA), and an air injector (Pneumatic MPPI-2 Pressure Injector, Eugene, OR). Injection mixes contained 1.30 μl of Cas9 enzyme (20 μM, New England BioLabs), 1.60 μl of prepared gRNAs, 2.5 μl of 4x Injection Buffer (0.2% phenol red, 800 mM KCl, 4 mM MgCl_2_, 4 mM TCEP, 120 mM HEPES, pH 7.0), and 4.6 μl of nuclease-free water.

### DNA isolation and genotyping

In order to determine the exact genetic alteration in our mutants, we performed Illumina amplicon sequencing through Massachusetts General Hospital DNA Core Facility (Cambridge, MA) using the primers found in Table S3. Briefly, a ~200 bp region surrounding the gRNA target site was amplified, purified using magnetic beads (AMPure XP, Beckman Coulter, Indianapolis, IN), and Illumina sequenced with 2 x 150 bp reads (Genewiz, San Diego, CA). Paired reads were aligned to the reference target region using *bwa* [50] using the zebrafish reference genome (GRCz11/danRer11). Specific alleles were determined using the R package CrisprVariants [51]. Genotyping stable lines was performed via PCR and SDS polyacrylamide gels or high-resolution melt curves (primers listed in Table S3). Adult zebrafish were anesthetized in 0.125% MS-222 and a small piece of caudal fin tissue (>50% of caudal fin) was dissected for crude DNA isolation. Briefly, 100 μl of 50 M NaOH was added to the fin clip and incubated in the solution at 95° for 20 min followed by 10 min 4° incubation. Then 10 μl of 5M HCL was added to the sample to neutralize the solution. The samples were vortexed for 5 seconds and the isolated DNA was used for PCR amplification and other downstream procedures. PCR amplification was performed using DreamTaq Green PCR master mix following the manufacturer’s protocol (Thermo Fisher Scientific, Waltham, MA) using the BioRad T100^™^ thermal cycler. High-resolution melt analysis was performed using SYBR green (Thermo Fisher Scientific) using the BioRad CFX96 Touch^™^ Real Time PCR System.

### Identification of potential CRISPR off-targets

CIRCLE-seq libraries for each gRNA and a control (Cas9 enzyme only) were prepared following the described protocol [31,52]. Illumina sequencing (Novogene, Sacramento, CA) of the libraries generated ~5 million reads per sample that were mapped to zebrafish reference genome (GRCz11/danRer11) to define predicted off-target sites relative to the control sample following the established CIRCLE-Seq bioinformatic pipeline. Once potential off-target sites were defined, we PCR amplified a ~500 bp region of G_0_ mosaic mutants of the top sites inside genes predicted by CIRCLE-seq (13 for *syngap1b* and 6 for *slc7a5*) (primers listed in Table S1) and ran a polyacrylamide gel electrophoresis to identify presence of indels at these sites by the formation of heteroduplexes [53]. Additionally, if heteroduplexes were detected, PCR reactions were cleaned-up using Ampure XP magnetic beads (Beckman Coulter), and Sanger sequenced (Genewiz, San Diego, CA). We did not observe off-target indels in these sites when compared to WT siblings.

### Statistical analysis

Statistical analyses were performed using R (3.5.0) for morphometric measurements and GraphPad Prism 8 software (GraphPad software, La Jolla, CA) for all behavioral assays. Pearson correlation tests were used to compare manual and automated measurements. Effect of factors on morphometric measurements was evaluated using multiple regression tests with the measurement as the response variable. PCA was performed using the R library PCAmixdata [54], which integrates PCA with multiple correspondence analyses (MCA) to evaluate quantitative (EA, L, Tel, OT) and qualitative (genotype) data simultaneously. Analysis of variance (ANOVA) was utilized to extract the effect of genotype in morphometric or behavioral measurements, followed by Tukey post-hoc tests to identify differences between genotypic groups. All models tested in this project controlled for inter-batch differences by adding ‘Experiment’ as a covariate. Kruskal-Wallis was used for behavioral analysis of mutants with a Dunn’s multiple comparison post-hoc tests performed for comparisons between genotypes. Significance from tests was defined by an alpha of 0.05.

## ACKNOWLEDGEMENTS

We thank our zebrafish husbandry personnel, including Bryant Palacios, Jenielee Mia, Andrew Nguyen, Sheena Fangonilo, and Breana Dyste for keeping our colony happy and healthy. We also thank Dr. Li-En Jao for his help and advice in the creation of our mutant lines, Bianca Yaghoobi for training using the Noldus DanioVision for behavioral assays, Chelsey Lee for contributions to off-target assessment of *slc7a5* mutants, and Daniela C. Soto for advice with bioinformatics. This work was funded in part by grants from the National Institutes of Health (NIH), including the National Institute of Neurological Disorders and Stroke (R00NS083627, M.Y.D.; U54 NS079202, P.J.L.), Office of the Director/National Institute of Mental Health (DP2 OD025824, M.Y.D.), and a UC Davis MIND Institute Intellectual and Developmental Disabilities Research Center pilot research grant funded through NIH National Institute of Child Health and Human Development (U54 HD079125, M.Y.D.). Additionally, M.Y.D. is supported as a Sloan fellow (FG-2016-6814), and A.Q. was supported through the RISE Program at Sacramento State University funded through the NIH (1R25GM122667). The funders played no role in the study design, data collection and analysis, decision to publish, or preparation of manuscript.

## SUPPORTING INFORMATION CAPTIONS

**Figure S1. Automated FishInspector quantification of morphometric features. (A)** Examples of zebrafish larvae at 3 and 5 dpf in dorsal and lateral view with FishInspector features highlighted. **(B)** Q-Q plots of each morphometric measurement (L, length, HW, head width; EA, average eye area; P, pericardium area; Y, yolk area) extracted from FishInspector with outliers in black with individuals highlighted purple if technical issues in imaging were observed. Plots include the best fit curve with 95% confidence intervals.

**Figure S2. Combined morphometric and behavioral experimental conditions tested.** WT NHGRI-1 larvae were subjected to different combinations of morphometric and behavioral tests across seven different experiments to determine their impacts on final phenotypic measurements. Not pictured are larvae that were tested with with a single experimental condition, including: behavioral measurements using the DanioVision at 5 dpf with 10 mM PTZ (n=52) and 0 mM PTZ (n=142) as well as morphometric measurements using the VAST at 3 dpf (n=185) and 5 dpf (n=61).

**Figure S3. Impact of morphometric and behavioral measurements by treatment variables.** Morphometric measurements of 5 dpf larvae subjected to varied treatments (VAST at 3 dpf, behavior at 5 dpf, and 10 mM of PTZ denoted with 0=no and 1=yes) as determined by the VAST system quantified via corrected FishInspector mappings for **(A)** L and **(B)** EA and via manual measurements for **(C)** Tel and **(D)** OT are shown as boxplots. **(E)** Average movement per minute quantified for the entire 1 h or the final 15 min after light flashing is shown as boxplots for larvae subjected to VAST measurements at 3 dpf (0=no and 1=yes) and at varied PTZ concentrations (red=0 mM and blue=10 mM). Total numbers of measured larvae are indicated in parentheses. Boxplots include the median value (dark line) and the whiskers represent a 1.5 interquartile range.

**Figure S4. Power to detect phenotypic differences from morphometric measurements using the combined platform.** For the four morphometric features measured at 5 dpf, we calculated how many samples are needed per group to achieve 80% power to detect varied percentages of changes between tested groups. For all traits, testing 50 larvae per group allows 4% change between measurements to be detected (shown by the dashed line).

**Figure S5. Generation and characterization of *syngap1b* CRISPR mutant. (A)** Phylogram indicates that two paralogs exist in zebrafish, *syngap1a* and *syngap1b*. Branch lengths were determined using a multiple-sequence alignment of CDS using the maximum-likelihood approach in MEGA7. **(B)** Longitudinal *syngap1b* expression in whole zebrafish embryos is plotted as transcript per million (TPM) using previously published data from White *et al*. (2017). **(C)** Top off-target sites hitting genes were identified for the gRNA used to generate the *syngap1b^tup1^* mutant using CIRCLE-seq. **(D)** Q-Q plots of each morphometric measurement with genotypes colored. Plotted on the x-axes are theoretical values from a normal distribution. Plots include the best fit curve with 95% confidence intervals. **(E)** Behavioral data from DanioVision motion tracking over 1 h without PTZ treatment, with genotypes colored as indicated. The arrow represents when flashing lights were administered for 1 min after 45 min of tracking. The plot represents the mean distance per minute with error bars represented as standard deviation.

**Figure S6. Generation and characterization of *slc7a5* CRISPR mutant. (A)** Phylogram indicates that one ortholog exists in zebrafish, *slc7a5*. Branch lengths were determined using a multiple-sequence alignment of CDS using the maximum-likelihood approach in MEGA7. **(B)** Longitudinal *slc7a5* expression in whole zebrafish embryos is plotted as transcript per million (TPM) using previously published data from White *et al*. (2017). **(C)** Top off-target sites hitting genes were identified for the gRNA used to generate the *slc7a5^tup1^* mutant using CIRCLE-seq. **(D)** Q-Q plots of each morphometric measurement with genotypes colored. Plotted on the x-axes are theoretical values from a normal distribution. Plots include the best fit curve with 95% confidence intervals. **(E)** Behavioral data from DanioVision motion tracking over 1 h without PTZ treatment, with genotypes colored as indicated. The arrow represents when flashing lights were administered for 1 min after 45 min of tracking. The plot represents the mean distance per minute with error bars representing standard deviation.

## REFERENCES

1. Meshalkina DA, N Kizlyk M, V Kysil E, Collier AD, Echevarria DJ, Abreu MS, et al. Zebrafish models of autism spectrum disorder. Exp Neurol. 2018;299: 207–216.

2. Grone BP, Baraban SC. Animal models in epilepsy research: legacies and new directions. Nat Neurosci. 2015;18: 339–343.

3. Sakai C, Ijaz S, Hoffman EJ. Zebrafish Models of Neurodevelopmental Disorders: Past, Present, and Future. Front Mol Neurosci. 2018;11: 294.

4. Howe K, Clark MD, Torroja CF, Torrance J, Berthelot C, Muffato M, et al. The zebrafish reference genome sequence and its relationship to the human genome. Nature. 2013;496: 498–503.

5. Flicek P, Amode MR, Barrell D, Beal K, Billis K, Brent S, et al. Ensembl 2014. Nucleic Acids Res. 2014;42: D749–55.

6. Kimmel CB, Ballard WW, Kimmel SR, Ullmann B, Schilling TF. Stages of embryonic development of the zebrafish. Dev Dyn. 1995;203: 253–310.

7. Jao L-E, Wente SR, Chen W. Efficient multiplex biallelic zebrafish genome editing using a CRISPR nuclease system. Proc Natl Acad Sci U S A. 2013;110: 13904–13909.

8. Chang N, Sun C, Gao L, Zhu D, Xu X, Zhu X, et al. Genome editing with RNA-guided Cas9 nuclease in zebrafish embryos. Cell Res. 2013;23: 465–472.

9. Peng Y, Clark KJ, Campbell JM, Panetta MR, Guo Y, Ekker SC. Making designer mutants in model organisms. Development. 2014;141: 4042–4054.

10. Wu RS, Lam II, Clay H, Duong DN, Deo RC, Coughlin SR. A Rapid Method for Directed Gene Knockout for Screening in G0 Zebrafish. Dev Cell. 2018;46: 112–125 e4.

11. Shah AN, Davey CF, Whitebirch AC, Miller AC, Moens CB. Rapid reverse genetic screening using CRISPR in zebrafish. Nat Methods. 2015;12: 535–540.

12. Varshney GK, Pei W, LaFave MC, Idol J, Xu L, Gallardo V, et al. High-throughput gene targeting and phenotyping in zebrafish using CRISPR/Cas9. Genome Res. 2015;25: 1030–1042.

13. Thyme SB, Pieper LM, Li EH, Pandey S, Wang Y, Morris NS, et al. Phenotypic Landscape of Schizophrenia-Associated Genes Defines Candidates and Their Shared Functions. Cell. 2019;177: 478–491.e20.

14. Pardo-Martin C, Chang T-Y, Koo BK, Gilleland CL, Wasserman SC, Yanik MF. High-throughput in vivo vertebrate screening. Nat Methods. 2010;7: 634–636.

15. Teixidó E, Kießling TR, Krupp E, Quevedo C, Muriana A, Scholz S. Automated Morphological Feature Assessment for Zebrafish Embryo Developmental Toxicity Screens. Toxicol Sci. 2019;167: 438–449.

16. Pulak R. Tools for automating the imaging of zebrafish larvae. Methods. 2016;96: 118–126.

17. Noldus LP, Spink AJ, Tegelenbosch RA. EthoVision: a versatile video tracking system for automation of behavioral experiments. Behav Res Methods Instrum Comput. 2001;33: 398–414.

18. LaFave MC, Varshney GK, Vemulapalli M, Mullikin JC, Burgess SM. A Defined Zebrafish Line for High-Throughput Genetics and Genomics: NHGRI-1. Genetics. 2014;198: 167–170.

19. Kozol RA, Cukier HN, Zou B, Mayo V, De Rubeis S, Cai G, et al. Two knockdown models of the autism genes SYNGAP1 and SHANK3 in zebrafish produce similar behavioral phenotypes associated with embryonic disruptions of brain morphogenesis. Hum Mol Genet. 2015;24: 4006–4023.

20. Hamdan FF, Daoud H, Piton A, Gauthier J, Dobrzeniecka S, Krebs M-O, et al. De novo SYNGAP1 mutations in nonsyndromic intellectual disability and autism. Biol Psychiatry. 2011;69: 898–901.

21. Tărlungeanu DC, Deliu E, Dotter CP, Kara M, Janiesch PC, Scalise M, et al. Impaired Amino Acid Transport at the Blood Brain Barrier Is a Cause of Autism Spectrum Disorder. Cell. 2016;167: 1481–1494.e18.

22. Näslund J. A simple non-invasive method for measuring gross brain size in small live fish with semi-transparent heads. PeerJ. 2014;2: e586.

23. Afrikanova T, Serruys A-SK, Buenafe OEM, Clinckers R, Smolders I, de Witte PAM, et al. Validation of the zebrafish pentylenetetrazol seizure model: locomotor versus electrographic responses to antiepileptic drugs. PLoS One. 2013;8: e54166.

24. Zheng Y-M, Chen B, Jiang J-D, Zhang J-P. Syntaxin 1B Mediates Berberine’s Roles in Epilepsy-Like Behavior in a Pentylenetetrazole-Induced Seizure Zebrafish Model. Front Mol Neurosci. 2018;11: 378.

25. Baraban SC, Taylor MR, Castro PA, Baier H. Pentylenetetrazole induced changes in zebrafish behavior, neural activity and c-fos expression. Neuroscience. 2005;131: 759–768.

26. Bandara SB, Carty DR, Singh V, Harvey DJ, Vasylieva N, Pressly B, et al. Susceptibility of larval zebrafish to the seizurogenic activity of GABA type A receptor antagonists. Neurotoxicology. 2020;76: 220–234.

27. Rumbaugh G, Adams JP, Kim JH, Huganir RL. SynGAP regulates synaptic strength and mitogen-activated protein kinases in cultured neurons. Proc Natl Acad Sci U S A. 2006;103: 4344–4351.

28. Walkup WG 4th, Washburn L, Sweredoski MJ, Carlisle HJ, Graham RL, Hess S, et al. Phosphorylation of synaptic GTPase-activating protein (synGAP) by Ca2+/calmodulin-dependent protein kinase II (CaMKII) and cyclin-dependent kinase 5 (CDK5) alters the ratio of its GAP activity toward Ras and Rap GTPases. J Biol Chem. 2015;290: 4908–4927.

29. Gamache TR, Araki Y, Huganir RL. Twenty Years of SynGAP Research: From Synapses to Cognition. J Neurosci. 2020;40: 1596–1605.

30. White RJ, Collins JE, Sealy IM, Wali N, Dooley CM, Digby Z, et al. A high-resolution mRNA expression time course of embryonic development in zebrafish. Elife. 2017;6. doi:10.7554/eLife.30860

31. Tsai SQ, Nguyen NT, Malagon-Lopez J, Topkar VV, Aryee MJ, Joung JK. CIRCLE-seq: a highly sensitive in vitro screen for genome-wide CRISPR-Cas9 nuclease off-targets. Nat Methods. 2017;14: 607–614.

32. Napolitano L, Scalise M, Galluccio M, Pochini L, Albanese LM, Indiveri C. LAT1 is the transport competent unit of the LAT1/CD98 heterodimeric amino acid transporter. Int J Biochem Cell Biol. 2015;67: 25–33.

33. Liu L, Yang G, Liu S, Wang L, Yang X, Qu H, et al. High-throughput imaging of zebrafish embryos using a linear-CCD-based flow imaging system. Biomed Opt Express. 2017;8: 5651–5662.

34. Reif DM, Truong L, Mandrell D, Marvel S, Zhang G, Tanguay RL. High-throughput characterization of chemical-associated embryonic behavioral changes predicts teratogenic outcomes. Arch Toxicol. 2016;90: 1459–1470.

35. Chang T-Y, Pardo-Martin C, Allalou A, Wählby C, Yanik MF. Fully automated cellular-resolution vertebrate screening platform with parallel animal processing. Lab Chip. 2012;12: 711–716.

36. Félix LM, Luzio A, Themudo M, Antunes L, Matos M, Coimbra AM, et al. MS-222 short exposure induces developmental and behavioural alterations in zebrafish embryos. Reprod Toxicol. 2018;81: 122–131.

37. Félix LM, Luzio A, Santos A, Antunes LM, Coimbra AM, Valentim AM. MS-222 induces biochemical and transcriptional changes related to oxidative stress, cell proliferation and apoptosis in zebrafish embryos. Comp Biochem Physiol C Toxicol Pharmacol. 2020;237: 108834.

38. Andropoulos DB. Effect of Anesthesia on the Developing Brain: Infant and Fetus. Fetal Diagn Ther. 2018;43: 1–11.

39. Ferreira LEN, Hasan D, Muniz BV, Sanchez JB, Volpato MC, Groppo FC. Effects of Local Anesthetics on Cellular Necrosis, Apoptosis and Inflammatory Modulation: Short Review. Journal of Anesthesia & Clinical Research. 2018. doi:10.4172/2155-6148.1000826

40. Topic Popovic N, Strunjak-Perovic I, Coz-Rakovac R, Barisic J, Jadan M, Persin Berakovic A, et al. Tricaine methane-sulfonate (MS-222) application in fish anaesthesia. J Appl Ichthyol. 2012;28: 553–564.

41. Kilinc M, Creson T, Rojas C, Aceti M, Ellegood J, Vaissiere T, et al. Species-conserved SYNGAP1 phenotypes associated with neurodevelopmental disorders. Mol Cell Neurosci. 2018;91: 140–150.

42. Parker MJ, Fryer AE, Shears DJ, Lachlan KL, McKee SA, Magee AC, et al. De novo, heterozygous, loss-of-function mutations in SYNGAP1 cause a syndromic form of intellectual disability. American Journal of Medical Genetics Part A. 2015. pp. 2231–2237. doi:10.1002/ajmg.a.37189

43. Galizia EC, Myers CT, Leu C, de Kovel CGF, Afrikanova T, Cordero-Maldonado ML, et al. CHD2 variants are a risk factor for photosensitivity in epilepsy. Brain. 2015;138: 1198–1207.

44. Warner TA, Liu Z, Macdonald RL, Kang J-Q. Heat induced temperature dysregulation and seizures in Dravet Syndrome/GEFS+ Gabrg2+/Q390X mice. Epilepsy Res. 2017;134: 1–8.

45. Peron A, Baratang NV, Canevini MP, Campeau PM, Vignoli A. Hot water epilepsy and SYN1 variants. Epilepsia. 2018. pp. 2162–2163.

46. Tegelenbosch RAJ, Lucas P J, Richardson MK, Ahmad F. Zebrafish embryos and larvae in behavioural assays. Behaviour. 2012;149: 1241–1281.

47. Wilson C. Aspects of larval rearing. ILAR J. 2012;53: 169–178.

48. Cassar S, Breidenbach L, Olson A, Huang X, Britton H, Woody C, et al. Measuring drug absorption improves interpretation of behavioral responses in a larval zebrafish locomotor assay for predicting seizure liability. J Pharmacol Toxicol Methods. 2017;88: 56–63.

49. Bonhomme V, Picq S, Gaucherel C, Claude J. Momocs: Outline Analysis UsingR. Journal of Statistical Software. 2014. doi:10.18637/jss.v056.i13

50. Li H, Durbin R. Fast and accurate short read alignment with Burrows-Wheeler transform. Bioinformatics. 2009;25: 1754–1760.

51. Lindsay H, Burger A, Biyong B, Felker A, Hess C, Zaugg J, et al. CrispRVariants charts the mutation spectrum of genome engineering experiments. Nat Biotechnol. 2016;34: 701–702.

52. Lazzarotto CR, Nguyen NT, Tang X, Malagon-Lopez J, Guo JA, Aryee MJ, et al. Defining CRISPR–Cas9 genome-wide nuclease activities with CIRCLE-seq. Nature Protocols. 2018. pp. 2615–2642. doi:10.1038/s41596-018-0055-0

53. Zhu X, Xu Y, Yu S, Lu L, Ding M, Cheng J, et al. An efficient genotyping method for genome-modified animals and human cells generated with CRISPR/Cas9 system. Sci Rep. 2014;4: 6420.

54. Chavent M, Kuentz-Simonet V, Labenne A, Saracco J. Multivariate Analysis of Mixed Data: The R Package PCAmixdata. arXiv [stat.CO]. 2014. Available: http://arxiv.org/abs/1411.4911

